# SARS-CoV-2 S1 spike protein induces a temporal systemic immune response and promotes long-term anxiety-like behaviors

**DOI:** 10.64898/2026.06.17.733010

**Authors:** Leyre Merino-Galan, John Hemenway, Hawa L. Jagana, Taylor S. Jackson, Ashmitha Rajendran, Asheema Khanna, Sergio Ortiz-Espinosa, Surojit Sarkar, Vandana Kalia, Siobhan S. Pattwell

**Affiliations:** Ben Towne Center for Childhood Cancer Research, Seattle Children’s Research Institute, Seattle, WA, United States of America; University of Washington School of Medicine, Seattle, WA, United States of America; Division of Human Biology, Fred Hutchinson Cancer Research Center, Seattle, WA, United States of America

**Keywords:** Long COVID, SARS-CoV-2, S1 protein, systemic immune response, cytokines, neurological symptoms, neuroinflammation, anxiety, amygdala

## Abstract

The SARS-CoV-2 S1 protein is associated with immune cell activation and persistent neurological symptoms, yet the underlying mechanisms remain unclear, posing a major challenge in elucidating Long COVID pathophysiology. To investigate how circulating S1 contributes to long-term neurological alterations, we intravenously injected hACE2 mice with varying doses of S1 (5, 10, and 20 µg) and observed temporally dysregulated systemic inflammatory responses accompanied by sub-acute CD4^+^ T cell infiltration into central limbic regions. This immune response induced mild sustained increases in cFos+ cells in the amygdala and mild neuroinflammation in the hippocampal CA1 region, resulting in both acute and long-term anxiety-like behaviors, while working memory remained unaffected. Together, these findings suggest that systemic S1 protein induces a sustained proinflammatory response that promotes lasting neurological alterations through immune-to-brain signaling pathways.

**Graphical Abstract:** 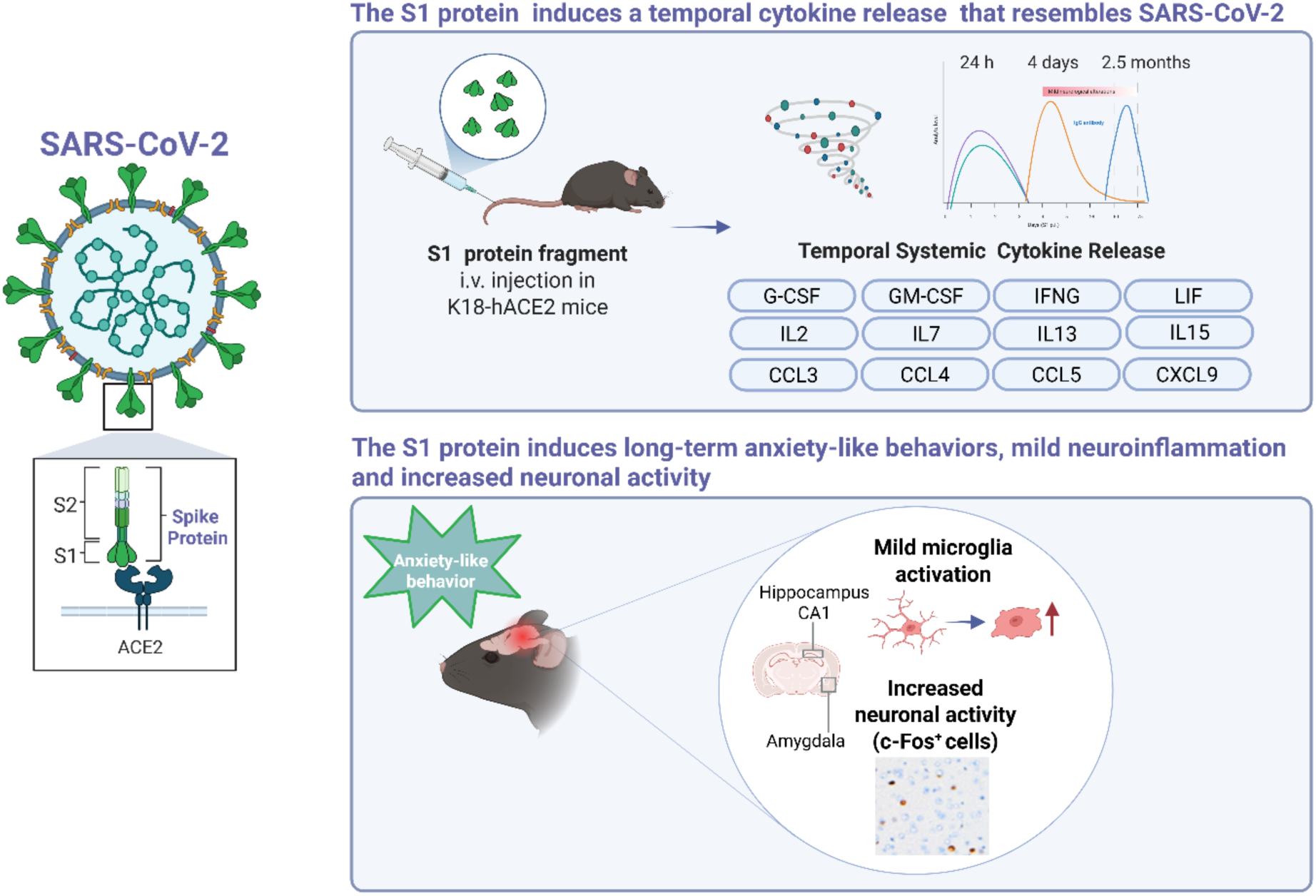

## Introduction

The **COVID-19** pandemic continues to have a profound impact on global health, with millions of cases reported worldwide (https://data.who.int), placing unprecedented strain on social, economic and health care systems. While the respiratory symptoms of SARS-CoV-2 infection are well-documented, its ability to affect multiple organ systems, including the brain, has also been recognized (Raman et al., 2021; Stein et al., 2022).

Although many individuals recover within weeks, a significant number experience post-acute sequelae of COVID-19 (PACS), commonly known as Long COVID, characterized by symptoms such as fatigue, headache, malaise and cognitive and neuropsychiatric disorders (Nasserie et al., 2021). Among these, neurological and psychiatric complications like attention deficits, anxiety and depression are particularly debilitating, affecting approximately 33.62% of COVID-19 patients after 6 months of diagnosis (Henry et al., 2020; Nasserie et al., 2021; Taquet, Geddes, et al., 2021; Taquet, Luciano, et al., 2021). Notably, these symptoms can persist even in individuals with mild initial COVID-19 symptoms (Becker et al., 2021; Nasserie et al., 2021; Vanichkachorn et al., 2021).

Despite strong clinical evidence for neurological symptoms in COVID-19 (Douaud et al., 2022), the underlying pathophysiological mechanisms remain unclear. Conflicting reports exist regarding the neuroinvasiveness of SARS-CoV-2, with some *post-mortem* studies showing direct SARS-CoV-2 infection and persistence of viral RNA in the brain (Matschke et al., 2020; Stein et al., 2022). Conversely, other studies failed to detect or found only minimal molecular traces of SARS-CoV-2 virus in the *post-mortem* brain tissue or cerebrospinal fluid (CSF), despite clear evidence of microglial activation and blood brain barrier (BBB) disruption (Greene et al., 2024; Schweitzer et al., 2022; Thakur et al., 2021; Yang et al., 2021). These findings suggest that mechanisms other than direct viral infection of brain parenchyma may be responsible for the observed neurological alterations. One hypothesis is that viral proteins, particularly the SARS-CoV-2 spike protein, may play a significant role (Rong et al., 2024).

The spike protein, composed of the S1 and S2 subunits, facilitates viral entry into host cells via the angiotensin-converting enzyme 2 (ACE2) receptor (Y. Huang et al., 2020). The S1 subunit binds to ACE2 and once cleaved during the viral entry process, can circulate in the bloodstream independently or in extracellular vesicles (Y. Huang et al., 2020; Troyer et al., 2021). Serum levels of the S1 protein have been associated with disease severity during the acute phase of COVID-19 (Ogata et al., 2020) and a recent study detected S1 protein in Long COVID patients 12 months post-diagnosis (Swank et al., 2023). In addition, it is known that the soluble S1 protein alone can induce systemic immune cell activation even in the absence of the virus itself (Chan et al., 2021; Henry et al., 2020) and it triggers neuroinflammation and neurological symptoms when directly inoculated into the brain (Erickson et al., 2023; Fontes-Dantas et al., 2023; Frank et al., 2022; Oh et al., 2022). However, direct brain inoculation of the S1 protein does not fully replicate the complex proinflammatory state observed in COVID-19. Reports of multisystem inflammatory syndrome (Feldstein et al., 2020; Riphagen et al., 2020) weeks after SARS-CoV-2 infection and neurological alterations following SARS-CoV-2 vaccination (Yang & Huang, 2023; Yonker et al., 2021) further underscore the potential impact of the spike protein on the nervous system.

However, the mechanisms by which the circulating SARS-CoV-2 S1 spike protein influences long-term neurological alterations and neuroinflammation are not fully understood, presenting a crucial challenge in addressing Long COVID. We hypothesized that circulating inflammatory S1 protein triggers a systemic proinflammatory response that, via immune-to-brain signaling pathways, leads to both acute and persistent central nervous system dysfunction. Accordingly, this study examines the temporal dynamics of systemic S1 administration, evaluating its effects on immune activation, neuroinflammation, and behavior to elucidate potential mechanisms linking peripheral inflammation to long-term neurological symptoms.

## Material and methods

### Animals and S1 protein administration

Adult C57BL/6J (000664) and K18-hACE2 mice (males and females, > 3 months old) were housed under standard conditions (70% humidity, 22°C, 12-h light/dark cycle) and with *ad libitum* access to food and water. C57BL/6J mice were obtained from Jackson Laboratory (Bar Harbor, ME, USA) and K18-hACE2 mice were bred in house.

A longitudinal study was performed on animals receiving either an intravenous (i.v.) injection of different doses (5, 10 or 20 μg) of the S1 protein (RayBiotech, #230-30161) or PBS. All the experimental procedures were approved by the local Institutional Animal Care and Use Committee at Seattle Children’s Research Institute (ACUC00698), and they were carried out in strict accordance with the US laws governing the treatment of animals. All efforts were made to minimize animal suffering and to reduce the number of animals used.

### Genotyping of hACE2 transgenic mice

Genomic DNA was isolated from mouse tail biopsies collected at the time of weaning using the standard alkaline lysis method with proteinase K. Two sets of screening were performed to detect the presence of the human angiotensin-converting enzyme 2 (hACE2) transgene – the first PCR was conducted to distinguish WT mice from transgenic mice on 3% agarose gel as per the JAX protocol <Protocol 38276 - Tg(K18-ACE2)2Prlmn-Chr2>, and then sanger sequencing assay PCR followed by restriction digestion by HpyCH4V enzyme was performed according to the JAX instructions <Protocol 38276 - Tg(K18-ACE2)2Prlmn-Chr2> to differentiate homozygous from heterozygous mice.

### Cytokine and chemokine analysis

To collect serum from mice, whole blood was obtained via chin bleeding, allowed to clot at room temperature, and then centrifuged to separate the serum. Mouse serum samples were diluted in DPBS (1:2) and serum cytokine and chemokine concentrations ̶ CCL11 (Eotaxin), G-CSF, GM-CSF, IFNG, IL1α, IL1β, IL2, IL3, IL4, IL5, IL6, IL7, IL9, IL10, IL12(p40), IL12(p70), IL13, IL15, IL17, CXCL10 (IP-10), CXCL1 (KC), LIF, CXCL5 (LIX), CCL2 (MCP-1), M-CSF, CXCL9 (MIG), CCL3 (MIP-1α), CCL4 (MIP-1β), CXCL3 (MIP-2), CCL5 (RANTES), TNFA, and VEGF ̶ were measured in duplicates using a Luminex Multiplex Assay (Bio-Plex 200, Bio-Rad). When cytokine levels were out of range (below the detectable threshold of the assay), they were set to 0 to account for undetectable or negligible concentrations and ensure consistency across the dataset.

### Neurophenotype Exam

Animals were habituated to being handled for several days before starting any experimental manipulation. An observer blinded to the treatment group conducted a daily 20-item neurophenotype exam (Faulhaber et al., 2022). Scored items include grooming, piloerection, respirations, extremity perfusion, eye-opening, body posture, tail posture, spontaneous activity, visual orienting, walk on cage edge, whisker reflex, eye blink, ear reflex, startle reflex, balance on the rod, climb onto the rod, reach for target from the suspended position, upper extremity grip strength, postural adjustment upon cage rotation and unusual behaviors. Each item was scored 0=normal, 1=performed with limitations/mildly abnormal or 2=unable to perform/severely abnormal, and the total daily score was determined by the sum of all items.

### Open Field Test

For *in vivo* monitoring of locomotor activity and anxiety-like behaviors, the open field test was carried out for 10 min in a 45 cm x 45 cm x 40 cm arena at baseline, 7 days, and 2 months p.i. Videos were acquired and analyzed with the SMART software (Panlab). The total distance travelled (cm), total velocity (cm/s), time spent in the central area (%) and immobility time (%) were analyzed. In addition, the experimenter manually quantified the fecal boli number at the end of the test. The arena was thoroughly cleaned with 70% ethanol and allowed to dry between trials.

### Novel Object Recognition Test

The novel object recognition test (NORT) was performed the next day following the open field test using the same arena and considering the open field test as the habituation phase for the NORT. In the NORT, during the familiarization phase, animals were exposed to two identical objects for 10 min. 24 h later, on the test phase, one of the objects was replaced by a new object (with different color and shape) and animals were allowed to explore the arena for 10 min. Videos were acquired and analyzed manually by an observer blinded to the treatment group. The time spent exploring each object during the familiarization and test phases were measured. Duration of exploration of objects during the familiarization phase provides a behavioral measure of attention (Alkam et al., 2011; Hornoiu et al., 2020). The discrimination index (DI) was calculated by subtracting the time spent exploring the familiar object from the time spent exploring the novel object, and then dividing this difference by the total exploration time of both objects. Given rodents’ innate preference for novelty, mice that successfully remember the familiar object will spend more time exploring the novel one, indicating intact recognition memory. To minimize olfactory cues between trials, the arena and objects were thoroughly cleaned with 70% ethanol and allowed to dry after each trial.

### Y-maze

Y-maze test was used to assess short-term spatial working memory. The mice were allowed to explore the arms freely over a period of 8 min. Videos were acquired and analyzed with the SMART software (Panlab) and the number of alternation triplets were analyzed. An alternation triplet is defined as consecutive entries into all three arms. This behavior is driven by the innate curiosity of rodents to explore previously unvisited areas. A high number of alternation triplets are taken as a good short-term spatial and working memory as this indicates that the mouse has recalled which arms it has already visited.

### Morris Water Maze

Morris Water Maze test was used to test spatial learning and memory. This test is based on two phases, the visible and the invisible phases, and a probe test. Mice underwent the visible-platform phase (eight trials per day) during three consecutive days, using a platform raised above the surface of the water and no visible cues. Then, mice undertook the hidden-platform phase where mice were trained during 5 consecutive days (four trials per day) to locate a platform submerged 1 cm beneath the water surface with the help of some visible cues present in the walls of the swimming pool. In this phase, non-toxic paint was used to make the water opaque, preventing the mice from seeing the platform. In both phases, mice were placed in selected locations, and each trial was finished either when the mouse reached the platform or after 60 s. After each trial, mice remained on the platform for 15 s. The next day after the hidden platform phase, mice were subjected to a probe trial (retention phase) in which they swam for 60 s in the pool without the platform. All trials were monitored by a camera connected to a SMART program (Panlab) for subsequent analysis of escape latencies during visible and hidden platform phases and percentage of time spent in each quadrant of the pool during the probe trial.

### Immunohistochemistry

Mice were euthanized with CO_2,_ and their brains were immediately collected. One hemisphere was immediately fixed in 10% neutral-buffered formalin for 72 h, then transferred to 70% ethanol for storage until further processing. Brains were embedded into paraffin blocks, sectioned in 10 μm serial slices, and mounted on slides. Immunohistochemical staining was performed using a Discovery Ultra Ventana Automated Stainer (Roche) with standard Ventana reagents (Roche). Brain sections were deparaffinated, subject to antigen retrieval (CC1, 950-124, Roche), and blocked to prevent non-specific staining. Sections were then incubated with the following primary and secondary antibodies: rabbit anti-Iba1 (1:200, 17198, Cell Signaling), rabbit anti-c-Fos (1:2000, ab222699, Abcam) and anti-rabbit HQ (760-4815, Roche). Sections were incubated with anti-HQ HRP (760-4820, Roche), developed using DAB (760-159, Roche), counterstained with hematoxylin (760-2021, Roche) and bluing reagent (760-2037, Roche) and coverslipped with mounting media.

### Image Analysis

Hippocampal and amygdala sections were imaged using the Thunder imager (Leica Biosystems, Germany) at 10x magnification. For each brain region, images were acquired from two consecutive sections and analyzed with FIJI software (NIH, https://fiji.sc/). For the hippocampus, 4–6 fields of view were obtained from the CA1, CA3 and *dentate gyrus* (DG) (348.8 × 348.8 µm) regions. In the amygdala, two images were captured, covering the lateral, basolateral and basomedial amygdala nuclei (669.7 × 669.7 µm). Microglia density and morphology were analyzed by a blinded experimenter, who manually counted and classified Iba-1^+^ cells into four morphometric types based on established criteria (Rodríguez-Chinchilla et al., 2020): ramified, dystrophic, bushy, and amoeboid. Briefly, the morphological characteristics of each type were defined as follows: Ramified microglia were cells with extended, thin processes and slight branching, with no detectable cytoplasm. Dystrophic microglia were cells with thicker processes, large soma, visible cytoplasm, and extensive branching. Bushy microglia were cells with irregular bodies, enlarged and less defined nuclei, shorter processes, and minimal branching. Amoeboid microglia were cells with large soma with few, thick, short processes; the nucleus occupies most of the cell body and is sometimes indistinguishable. cFos^+^ cells were quantified from DAB-extracted images using a custom macro in FIJI. Images were first converted into binary masks by applying a threshold to distinguish foreground objects from the background. cFos^+^ nuclei were identified based on predefined criteria: only particles larger than 40 pixels were considered, and a circularity threshold of 0.30–1.00 was applied. cFos^+^ cells were visualized as masks, and the number of cFos^+^ nuclei per image was automatically quantified. All imaging and quantification were performed by a researcher blinded to the experimental conditions. Data were grouped by animal and experimental condition.

### Statistical Analysis

Statistical analyses were performed using GraphPad Prism 10.3.1 (GraphPad Software Inc.). Normality of data distribution was assessed with Shapiro-Wilk test. For multiple comparisons between the S1 and PBS groups, one-way ANOVA followed by Dunnet’s *post hoc* test was used. In cases involving repeated measures or two independent factors, two-way ANOVA followed by Dunnet’s *post hoc* test was applied. Correlation analyses were performed using Pearson’s correlation coefficient. Data are represented as mean ± SEM, with significance set at *P* < 0.05.

## Results

### SARS-CoV-2 spike S1 protein triggers temporal and dose-dependent systemic inflammation characterized by an elevated cytokine release in K18-hACE2 mice

To determine if the circulating SARS-CoV-2 spike S1 protein can induce systemic immune activation *in vivo*, K18-hACE2 and WT mice were intravenously injected with different doses of the S1 protein, and serum levels of 32 cytokines and chemokines were measured over time (**Figure 1B**). The S1-injected K18-hACE2 mice exhibited a dose-dependent and time-dependent increase in cytokine release (**Figure 1C, D, E**), whereas no significant cytokine release was observed in WT mice (**Figure S1**), indicating that the cytokine release observed is dependent on hACE2 expression in circulating immune cells or vascular cells.

**Figure 1.**
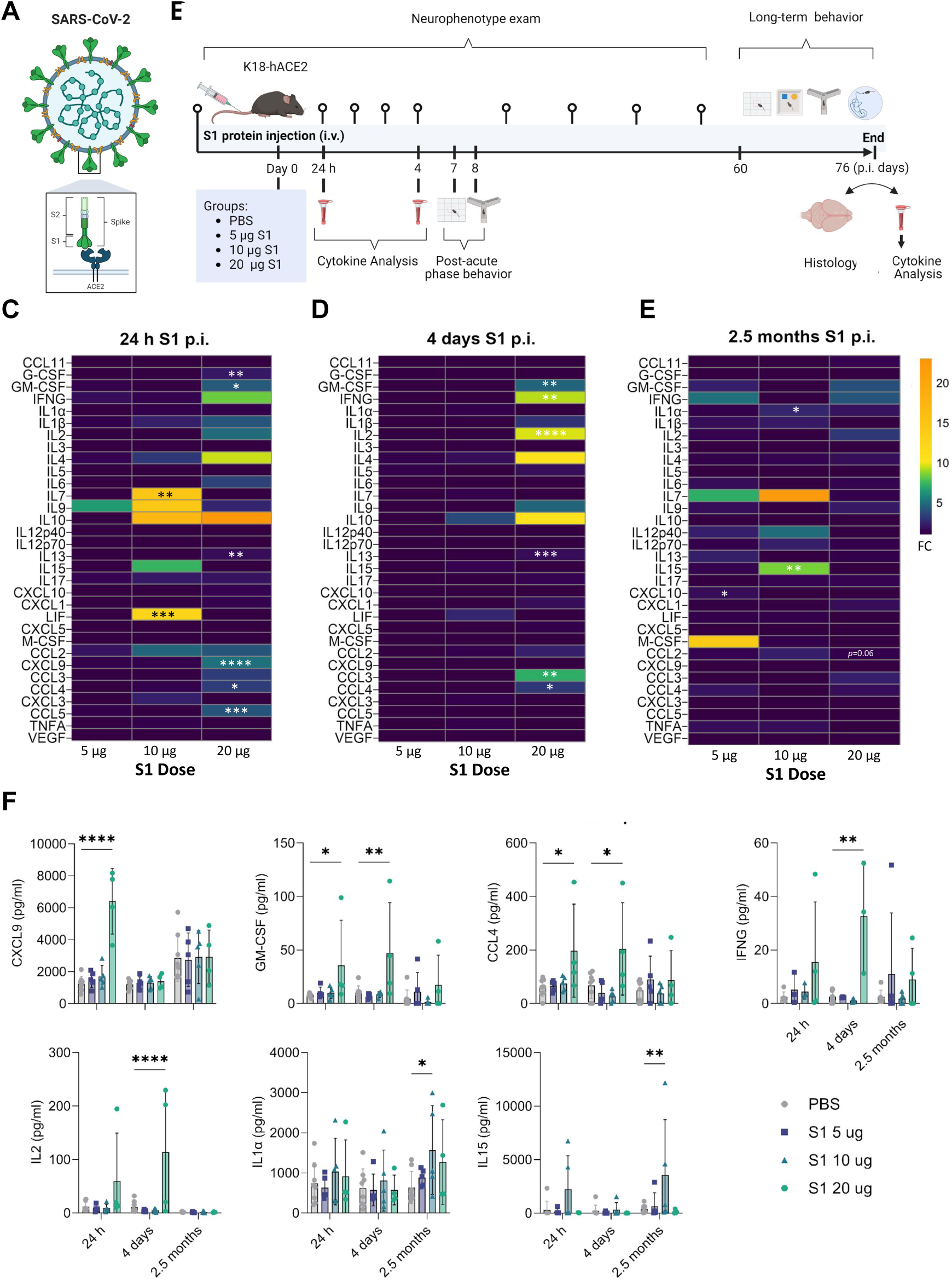
SARS-CoV-2 spike S1 protein triggers temporal and dose-dependent cytokine and chemokine elevations in K18-hACE2 mice. (**A**) Schematic representation of SARS-CoV-2 virus and Spike protein. (**B**) Schematic of experimental paradigm for SARS-CoV-2 S1 protein injection (intravenous, i.v) in K18-hACE2 mice and experimental workflow. (**C**, **D** and **E**) Cytokine analyses of serum in control (PBS) and S1-injected mice at different doses (Created with biorender.com.) at 24 h (C), 4 days (D) and 2.5 months post-inoculation (p.i) (E). Data shown as fold change (FC) of the concentration corresponding to the median fluorescence intensity compared with control group; n = 4–7 mice per group. (**F**) Cytokine analyses of serum in control and S1-injected mice at different doses (5 µg, 10 µg and 20 µg) 24 h, 4 days and 2.5 months post-inoculation (p.i.) Data shown as the concentration (pg/ml) corresponding to the median fluorescence intensity. n = 4–7 mice per group. Illustrations were created with biorender.com. Data are presented as mean ± SEM. Each dot represents an individual mouse. Repeated measure Two-way ANOVA test followed by Dunnet’s *post-hoc* test: \**P* < 0.05; \*\**P* < 0.01, \*\*\**P* < 0.001, \*\*\**P* < 0.0001.

At 24 h p.i., elevated levels of CXCL9, G-CSF, GM-CSF, IL13, CCL4, CCL5, IL7, LIF were detected in the serum of S1-injected mice (**Figure 1C**). Notably, GM-CSF, CCL4, IL13 remained elevated at 4 days p.i. in the group treated with 20 µg, with additional increase in IFNG and IL2 and CCL3 observed at this time point and group (**Figure 1D**). While the initial inflammatory response diminished over time, residual inflammation persisted. At 2.5 months p.i., IL1α and IL15 levels were still elevated in the 10 µg S1 group and CXCL10 in the 5 µg S1 group compared to the control group (**Figure 1E**).

However, our serum profiling did not reveal significant changes in markers of vascular inflammation or endothelial dysfunction following seven days after S1 administration (**Figure S2**). Although we observed a trend toward decreased sE-Selectin and increased sP-Selectin in several mice from the 20 μg S1 group compared to PBS, these differences were not statistically significant. This suggests the absence of sustained or systemic endothelial activation in this model, or possibly reflects a transient or resolving inflammatory phase, as samples were collected during the sub-acute stage (7 days p.i.).

Overall, these findings show that K18-hACE2 mice develop an acute and transient cytokine release following the intravenous injection of the S1 protein, which is dependent on the hACE2 expression and mirrors the inflammatory response observed in COVID-19 patients.

### SARS-CoV-2 spike S1 protein induces anxiety-like behaviors and attention deficits without memory alterations in K18-hACE2 mice

To investigate whether the systemic S1 protein contributes to the neurological symptoms observed in COVID-19 patients, K18-hACE2 mice were monitored for two months to detect signs of neurological abnormalities and behavioral changes. Mice exhibited subtle neurological abnormalities (**Figure 2A**) without weight loss (**Figure 2B**). Administration of 20 μg of S1 protein led to a mild increase in neuro score from 4 days p.i. onwards compared to baseline (**Figure 2A, Figure S2**), while the 10 µg S1 group showed an increase in neuro score at 44 days p.i., though this did not reach statistical significance (p= 0.07) (**Figure 2A)**. The most frequent abnormalities on the neuro score were deficits in postural adjustment and balance. No significant differences in the neuro score were noted among WT mice between groups (**Figure S3A**).

**Figure 2.**
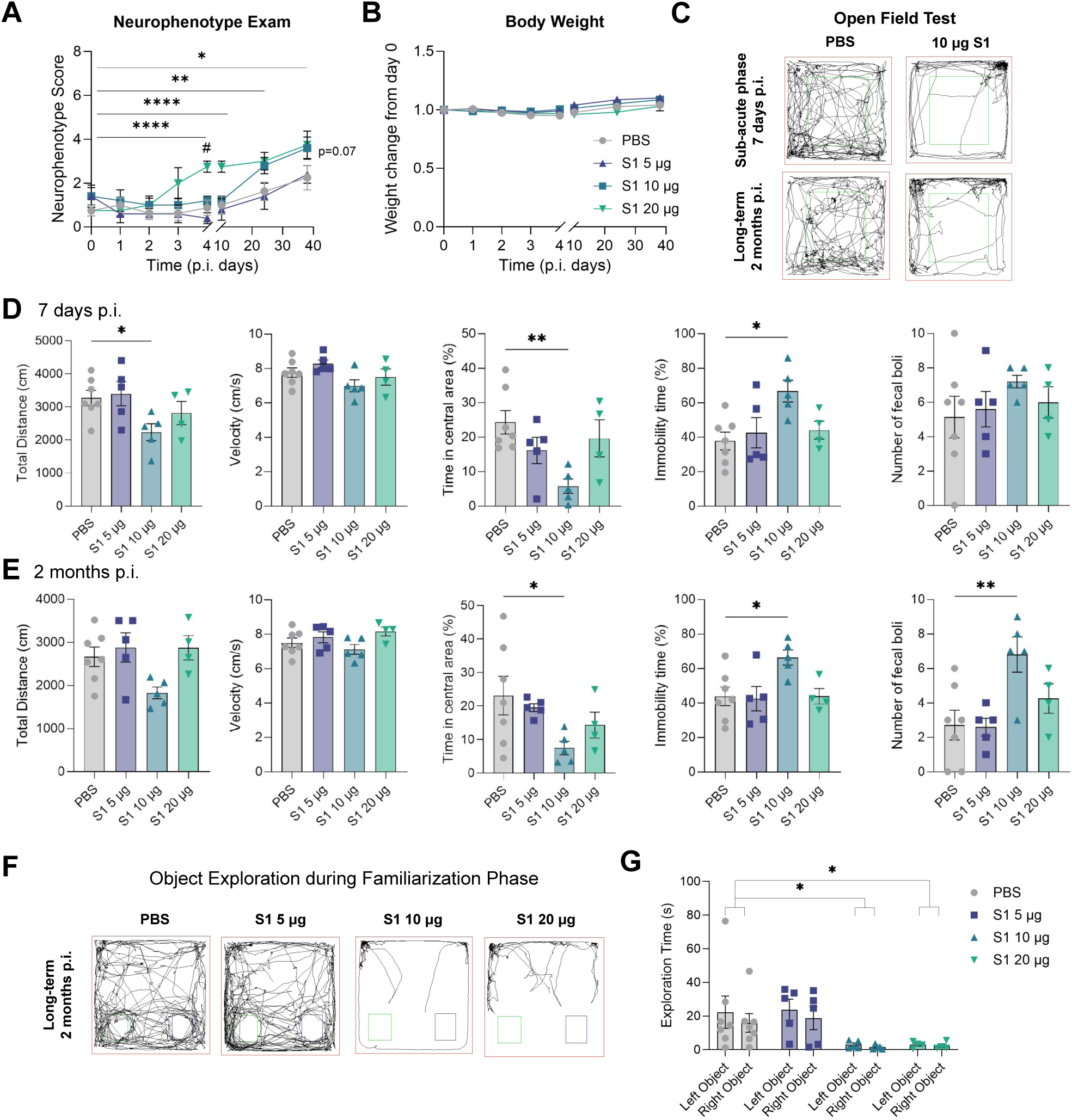
SARS-CoV-2 spike S1 protein induces anxiety-like behaviors and exploration deficits. (**A**) Neurophenotype scores over time in control and S1-injected mice at different doses (5 µg, 10 µg and 20 µg). Repeated measure Two-way ANOVA followed by Dunnet’s *post-hoc* test: ns, *P* > 0.05, \**P* < 0.05; \*\**P* < 0.01, \*\*\**P* < 0.001, \*\*\**P* < 0.0001 vs. time 0. ^#^*P* < 0.05 vs. PBS. (**B**) Body weight (change from day 0) over time in control and S1-injected mice at different doses (5 µg, 10 µg and 20 µg). (**C**) Open field test representative track plots with traces of mouse movement. (**D and E**) Total distance travelled (cm), velocity (cm/s), time in central area (%), immobility time (%) and number of fecal boli in control and S1-injected mice at different doses (5 µg, 10 µg and 20 µg) 7 days p.i. (D) and 2 months p.i. (E). One-way ANOVA followed by Dunnet’s *post-hoc* test: \**P* < 0.05; \*\**P* < 0.01. (**F**) Representative track plots with traces of mouse object exploration during the familiarization phase of the novel object recognition test. (**G**) Exploration time (s) of two identical objects (left and right) during the familiarization phase of the novel object recognition test in control and S1-injected mice at different doses (5 µg, 10 µg and 20 µg) 2 months p.i.. Two-way ANOVA followed by Dunnet’s *post-hoc* test: \**P* < 0.05. Data shown as mean ± SEM. Each dot represents an individual mouse. n = 4-7 mice per group.

In the open field test, mice injected with the 10 µg dose of S1 spent less time in the central area and exhibited increased immobility compared to the control group at both 7 days and at 2 months p.i. (**Figure 2C-E**), indicative of anxiety-like behavior.

Furthermore, the number of fecal boli, another indicator of anxiety, was significantly elevated in the 10 µg group at 2 months p.i. (**Figure 2E**). Locomotor function, assessed by the total distance traveled and mean velocity, was significantly altered only at 7 days p.i. (total distance) in the 10 µg group (**Figure 2D**). However, no significant changes in these parameters were observed in the Y-maze test (**Figure 3B**) or in the neurophenotype exam, suggesting that the reduced activity in the open field test likely reflects anxiety-like behavior than locomotor impairments. Object exploration during the familiarization phase revealed a significant reduction in exploration time in both the 10 µg and 20 µg groups compared to the control group at 2 months p.i. (**Figure 2F, G),** indicative of anxiety-like or attention deficits. This decrease in exploration made it difficult to obtain reliable results in the subsequent test phase of the NORT, as insufficient familiarization with the objects likely impaired the animals’ ability to form a memory of them (**Figure S5**).

**Figure 3.**
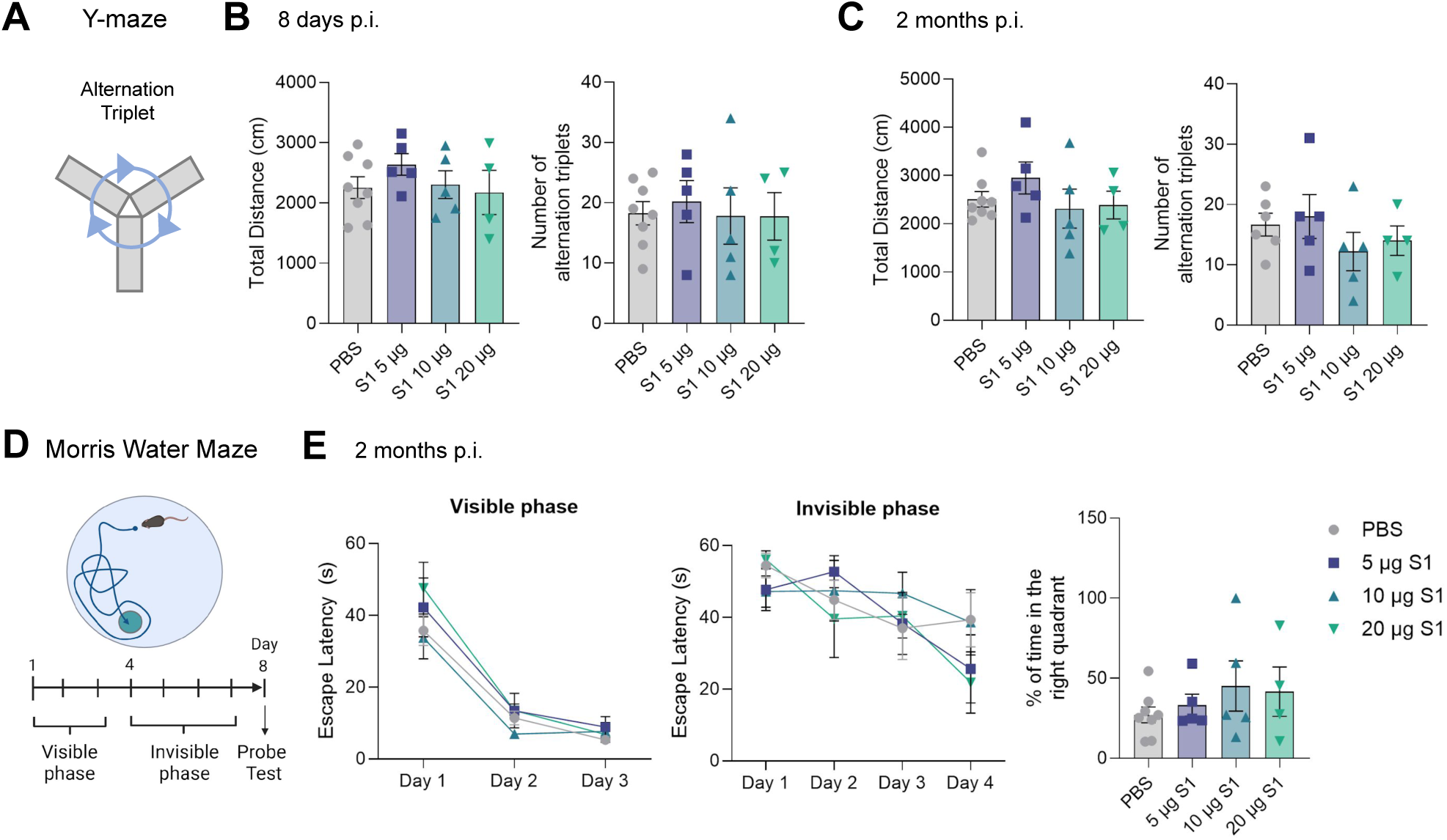
SARS-CoV-2 spike S1 protein is not associated with memory alterations in K18-hACE2 mice. **(A)** Schematic representation of the Y-maze. **(B, C)** Total distance (cm) and number of alternation triplets in control and S1-injected mice at different doses (5 µg, 10 µg and 20 µg) 7 days p.i. (B) and 2 months p.i. (C) during the Y-maze test. One-way ANOVA: no significant differences observed. (**D**) Schematic representation of the Morris water maze test. (**E**) Escape latency (s) during the visible and invisible phases and % of time spent in the right quadrant during the probe test of the Morris water maze, in control and S1-injected mice at different doses (5 µg, 10 µg and 20 µg) 2 months p.i. Repeated measure two-way ANOVA and one-way ANOVA, respectively: no significant differences observed. Data shown as mean ± SEM. Each dot represents an individual mouse. n = 4-7 mice per group.

We also evaluated the learning and spatial memory of these mice. In the Y-maze test, neither S1 protein-injected group displayed altered short-term spatial memory at either 8 days or 2 months p.i. compared to the control group as shown by the number of alternation triplets (**Figure 3A, B, C**). Similarly, in the Morris water maze test, no significant differences in learning during the visible phase or spatial working memory during the invisible phase and the test day were observed between the S1-injected and control groups at 2 months p.i. (**Figure 3D, E**). No significant differences in the open field test, Y-maze, NORT or Morris water maze were noted among WT mice between groups (**Fig S3C, D, E, F**).

Taken together, these results indicate that the intravenous administration of SARS-CoV-2 S1 protein induced significant anxiety-like behavior and attention deficits in K18-hACE2 mice but not in WT mice, both in the sub-acute and long-term phases, without impairing locomotor activity, learning or spatial working memory.

### The release of specific cytokines following S1 administration correlates positively with the neurophenotype score during the acute phase, though only a few cytokines maintain a significant correlation with behavioral parameters in the long-term

To determine whether cytokine release is directly linked to behavioral changes, we performed a correlation analysis. A positive correlation was observed between the levels of CCL11, IL2, IL13, CXCL10 and CXCL9 levels at 24 h p.i. and the neurophenotype score at day 4 p.i. (**Figure 4A, D**). However, no significant correlations were found between cytokine levels at 24 h and the long-term progression of the neurophenotype score (**Figure S6C**). Similarly, at 4 days p.i., a positive correlation emerged between GM-CSF, IFNG, IL2, IL9, CXCL1 and CCL3 levels and the neurophenotype score at 4 days p.i., though only IL3 showed a significant negative correlation with the neurophenotype score in the long-term (**Figure S6A, D**).

**Figure 4.**
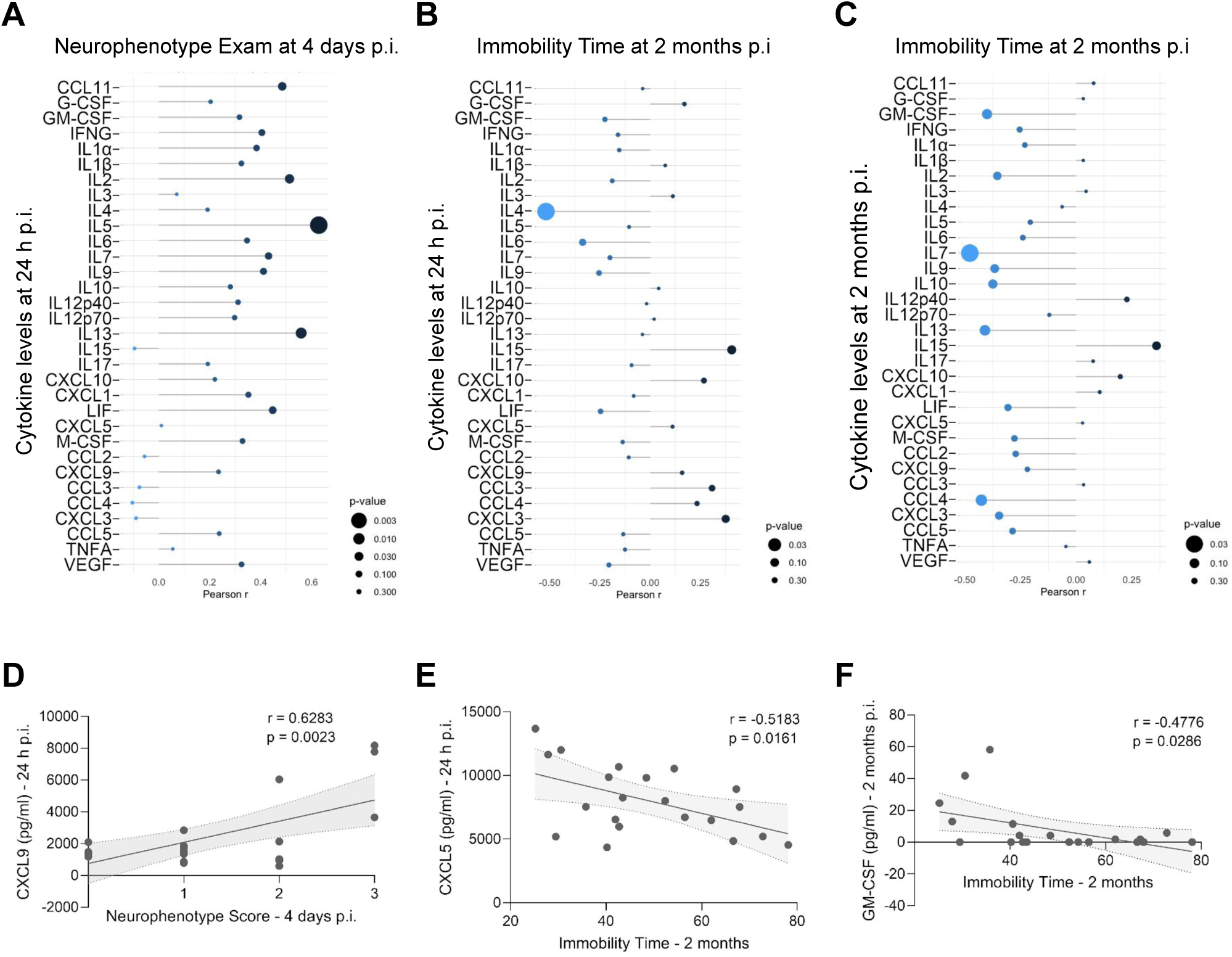
Correlations of cytokine levels with behavioral outcomes during the acute and long-term phases. (**A** and **B**) Correlation of the cytokine levels at 24 h p.i. with the neurophenotype exam at 4 days p.i. (**A**) and immobility time at 2 months p.i. (**B**). (**C**) Correlation of the cytokine levels and immobility time at 2 months p.i. (**D**) Positive correlation of CXCL9 levels at 24 h p.i. with neurophenotype score at 4 days p.i. (**E**) Negative correlation of CXCL5 levels at 24 h p.i. with immobility time at 2 months p.i. (**F**) Negative correlation of GM-CSF levels with immobility time at 2 months p.i. Pearson’s correlations. Total n = 21 mice.

When examining the correlation between the cytokine levels and long-term anxiety-like behaviors, we found that at 24 h p.i. only CXCL5 showed a significant negative correlation with long-term immobility time (**Figure 4B, E**), while no significant correlations were observed at 4 days post-infection (**Figure S6B**). At 2 months post-infection, only GM-CSF levels showed a negative correlation with immobility time (**Figure 4C, F**). These findings suggest that cytokine levels during the acute phase are more closely associated with the immediate neurological state than with long-term behavioral outcomes.

### Systemic administration of SARS-CoV-2 spike S1 protein induces persistent mild microglial activation in the hippocampus and neuronal activation in the amygdala

To investigate potential pathological changes in the brains of mice injected with the S1 protein, we focused on evaluating signs of neuroinflammation in key regions of the limbic system involved in fear processing and anxiety, specifically the hippocampus and amygdala. Although no significant changes were observed in the total number of Iba1^+^ cells across the analyzed regions (**Figure 5A, B**), we detected an increased number of ameboid-shaped Iba1^+^ microglia in the CA1 region of the hippocampus in the 10 μg S1 group (**Figure 5C, D**), indicating the activation and persistence of mild neuroinflammatory processes. While the hippocampal CA1, CA3, and DG regions are all implicated in anxiety regulation, the ventral CA1 subregion plays a particularly prominent role due to its direct connectivity with anxiety-related structures such as the amygdala and hypothalamus (Sokolowski & Corbin, 2012). To evaluate neuronal activation, we quantified cFos^+^ cells and observed a significant increase in the amygdala of mice treated with 10 μg of S1 (**Figure 5F, G**). This finding, aligned with the group exhibiting pronounced anxiety-like behaviors, suggests that peripheral exposure to the S1 protein can trigger long-lasting alterations in limbic circuit activity associated with emotional processing.

**Figure 5.**
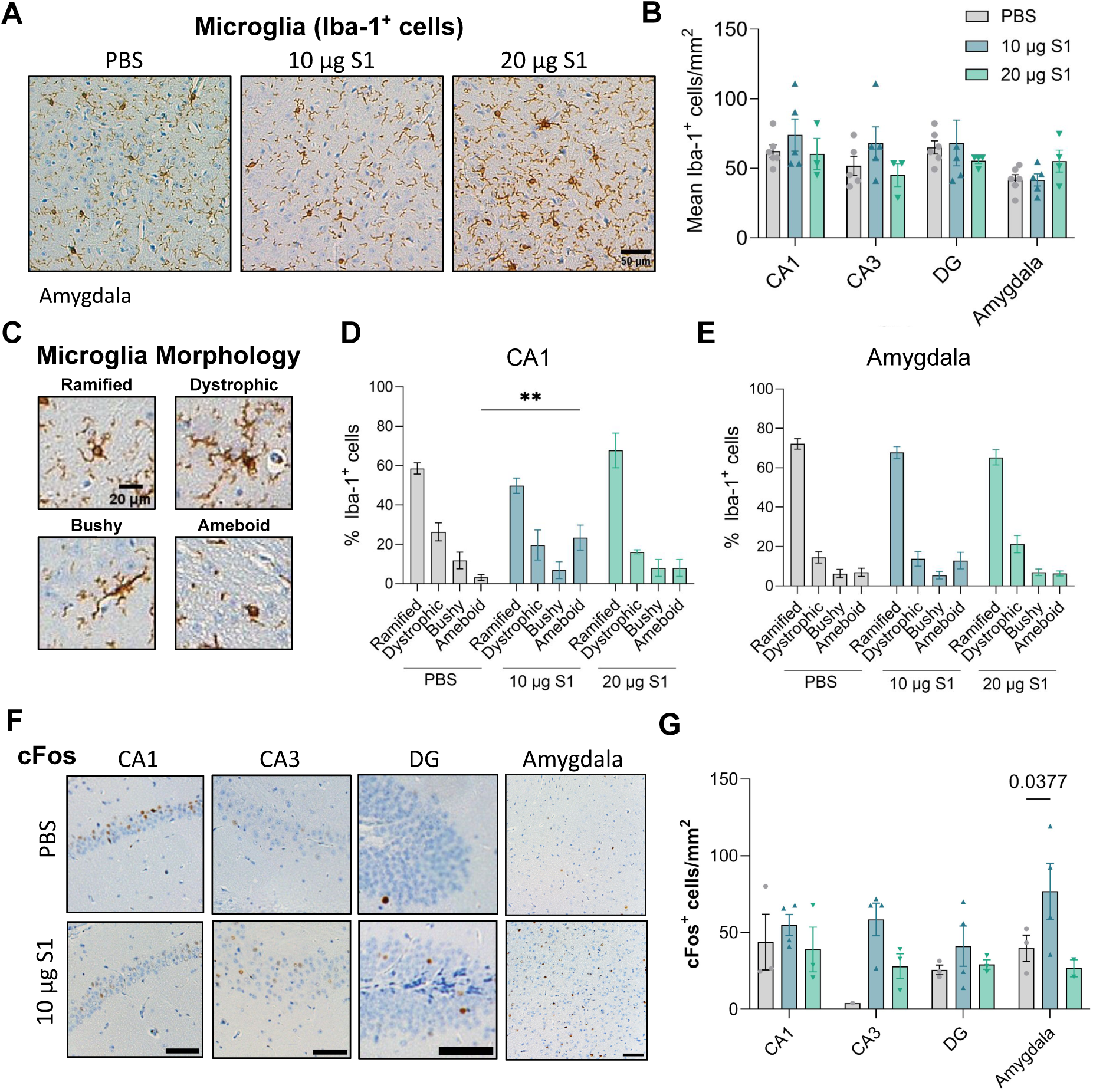
SARS-CoV-2 S1 protein induces sustained mild neuroinflammation in the hippocampus and neuronal activation in the amygdala. **(A)** Representative images of Iba-1^+^ cells in the amygdala in control and S1-injected mice at different doses (10 µg and 20 µg) 2.5 months p.i. Scale bar, 50 µm. **(B)** Mean Iba-1^+^ cells per mm^2^ in the hippocampus regions (CA1, CA3 and DG) and amygdala of control and S1-injected mice at different doses (10 µg and 20 µg) 2.5 months p.i. **(C)** Representative images of Iba-1^+^ microglia displaying distinct morphologies: ramified, dystrophic, bushy, and amoeboid. Scale bar, 20 µm. **(D, E)** Percentage of Iba-1^+^ microglial morphology types in the CA1 region (**D**) and amygdala (**E**) of control and S1-injected mice at different doses (10 µg and 20 µg) 2.5 months p.i (**F**) Representative images of cFos^+^ cells in the hippocampus regions (CA1, CA3, DG) and amygdala of control and S1-injected mice with 10 µg dose at 2.5 months p.i. Scale bars, 100 μm. (**G**) Mean cFos^+^ cells per mm^2^ in the hippocampus regions (CA1, CA3 and DG) and amygdala of control and S1-injected mice at different doses (10 µg and 20 µg) 2.5 months p.i. Data shown as mean ± SEM. Each dot represents an individual mouse. n = 3-4 mice per group. Two-way ANOVA followed by Dunnet’s *post-hoc* test: \**P* < 0.05, \*\**P* < 0.01.

### Systemic S1 exposure induces sub-acute CD4^+^ T-cell accumulation in limbic brain regions

Although no signs of endothelial damage were detected during the sub-acute stage (**Figure S2C**), we investigated whether systemic S1 protein exposure could trigger immune cell infiltration into key limbic regions 8 days p.i. — including core limbic structures (amygdala and hypothalamus), the hippocampus, and the prefrontal cortex (**Figure S2A**) —which might contribute to the long-term histopathological alterations observed. We detected a trend toward increased CD45^+^ cell frequency in the core limbic structures and the hippocampus in the 20 μg S1 group compared with PBS controls (**Figure S7A**) along with a significant increase in CD4^+^ T cells in the central limbic regions in the 20 μg S1 group (**Figure S7B**). Together, these findings suggest that systemic exposure to S1 protein may promote low-grade immune cell infiltration into limbic brain regions, potentially contributing to the neuroinflammatory processes underlying the long-term neuropathological changes observed in this model.

## Discussion

Here we demonstrate that systemic exposure to the SARS-CoV-2 spike S1 protein is sufficient to induce a temporally dysregulated inflammatory response that is dependent on hACE2 expression and associated with both sub-acute and persistent anxiety-like behaviors. These behavioral alterations co-occur with mild but sustained neuroinflammation in the hippocampal CA1 region and prolonged increase in cFos+ cells in the amygdala, supporting a peripheral immune-to-brain mechanism contributing to long-term neuropsychiatric sequelae relevant to Long COVID.

Circulating S1 protein has been detected during acute COVID-19 (Ogata et al., 2020) and up to 12 months post-infection in individuals with Long COVID (Swank et al., 2023), but not in recovered controls, suggesting prolonged antigenic exposure may occur independently of productive viral replication. Viral proteins from other pathogens, such as HIV gp120, are known to exert neuroinflammation and neurocognitive deficits in the absence of active infection (Kesby et al., 2015; Lee et al., 2021), supporting the concept that S1 may function as an immune-modulating factor rather than merely a structural viral component. Consistent with prior *in vitro* and *in vivo* studies (Chan et al., 2021; Olajide et al., 2021; Shirato & Kizaki, 2021), we show that S1 alone triggers robust immune activation, reinforcing its capacity to drive pathology independently of whole virus.

Using a broad cytokine panel, we observed that S1 administration elicited an acute cytokine profile partially overlapping with that reported in severe COVID-19 (Blanco-Melo et al., 2020; Costela-Ruiz et al., 2020; Del Valle et al., 2020; C. Huang et al., 2020; Merad & Martin, 2020; Olbei et al., 2021), including increases in GM-CSF, CXCL9, CXCL10, IFNG, IL2, and IL13. These mediators implicate both innate and adaptive immune pathways and mirror cytokine signatures linked to neuropsychiatric manifestations in COVID-19 and Long COVID patients (Greene et al., 2024; Phetsouphanh et al., 2022). Although the inflammatory response diminished over time, persistent elevation of IL1α, IL15, and CXCL10 two months after a single S1 exposure suggests long-lasting immune reprogramming (Becher et al., 2017). The differences in cytokine responses between doses may be attributed to the pharmacodynamics of the S1 protein, potentially influencing both the kinetics and magnitude of cytokine release. However, the contribution of biological and experimental variability cannot be excluded. Notably, acute cytokine levels correlated with early neurological abnormalities but only weakly predicted long-term behavioral outcomes, supporting the notion that transient inflammation may initiate downstream processes that become independent of ongoing cytokine elevation.

Behaviorally, S1-injected hACE2 mice exhibited sustained anxiety-like behaviors and reduced object exploration without impairments in learning or spatial memory. This phenotype parallels clinical observations in Long COVID, where anxiety and attentional deficits are prevalent neuropsychiatric symptoms (Nasserie et al., 2021). Research in both humans and animals has demonstrated that elevated levels of circulating cytokines can lead to “;sickness behaviors,” characterized by reduced general activity, diminished exploration, anhedonia, and disruptions in sleep and learning (Dantzer, 2001; Hou et al., 2017), as well as, neuropsychiatric disorders (Dantzer et al., 2008; Kronfol & Remick, 2000). Typically, these sickness behaviors resolve once the acute phase of the illness subsides. However, exacerbated neuroinflammation can result in extended sickness behaviors, cognitive impairments, and increased sensitivity to stress (Godbout et al., 2005). At the neurobiological level, S1 exposure resulted in increased ameboid microglia specifically in the hippocampal CA1 region and enhanced cFos expression in the amygdala – regions critically involved in anxiety regulation and stress responsivity. The ventral CA1 – amygdala circuit is particularly sensitive to inflammatory modulation, and persistent dysregulation within this network may underlie the behavioral abnormalities observed.

While leukocyte trafficking into the CNS is tightly regulated under physiological conditions, we observed low-grade immune cell infiltration, particularly of CD4^+^ T cells, within limbic regions during the sub-acute phase. Although no overt endothelial dysfunction was detected at the assessed time point, immune cell entry may have been facilitated by transient BBB alterations that resolved before analysis. In addition, peripheral cytokines can signal across the BBB and activate resident glial cells, amplifying neuroinflammatory responses even in the absence of persistent barrier disruption (Becher et al., 2017; Dantzer, 2018). Together, these mechanisms may contribute to sustained changes in CNS immune homeostasis and neuronal activity.

An alternative, non-mutually exclusive mechanism involves direct S1 entry into the brain. The S1 protein has been shown to cross the BBB (Erickson et al., 2023; Rhea et al., 2021), and human ACE2 is highly expressed in limbic regions, including the amygdala and brainstem (Lukiw et al., 2022). Prior studies using direct intracerebral S1 administration demonstrated strong neuroinflammatory and behavioral effects (Erickson et al., 2023; Fontes-Dantas et al., 2023; Frank et al., 2022; Oh et al., 2022) but lacked the peripheral inflammatory context of COVID-19 and employed WT mice with murine ACE2, which exhibits reduced compatibility and lower binding efficiency with SARS-CoV-2 spike protein (Lan et al., 2020; Zhou et al., 2020). Our hACE2-dependent model may therefore better capture the combined peripheral and central actions of S1 relevant to human disease.

Persistent viral reservoirs of SARS-CoV-2 releasing S1 protein to the bloodstream have been proposed as a contributor to Long COVID pathogenesis (Proal et al., 2023; Rong et al., 2024). Although our study employed a single systemic exposure, the sustained behavioral and neuroimmune changes observed suggest that even transient inflammatory insults may be sufficient to induce enduring CNS dysfunction (Dantzer, 2001). The persistence of single-stranded RNA viruses, such as Ebola (Keita et al., 2019) and Zika (Paz-Bailey et al., 2018), long after the acute phase of illness provides precedent for this phenomenon. Chronic or repeated S1 exposure, potentially mimicking viral persistence, may exacerbate these effects and warrants further investigation.

Several limitations should be acknowledged. The study was not powered for sex-specific analyses, and cerebrospinal fluid was not collected to directly assess BBB integrity. Additionally, only the ancestral S1 protein was examined and emerging variants with altered receptor affinity or immunogenicity may exhibit distinct neuroimmune effects.

In summary, our findings identify circulating SARS-CoV-2 S1 protein as a potent driver of systemic immune activation capable of producing lasting neuroimmune and behavioral alterations. These results provide mechanistic insight into how peripheral viral proteins may contribute to long-term neuropsychiatric symptoms in COVID-19, supporting immune-to-brain signaling as a central component of Long COVID pathophysiology.

## Acknowledgements

We would like to thank Ginny Schultz from the Histology Core at Seattle Children’s Research Institute (SCRI) for her technical assistance on immunohistochemistry. We also thank SCRI for the use of their Core Facilities and infrastructure, as well as the Flow Cytometry Core at Fred Hutchinson Cancer Institute for their continuous support. We acknowledge Creative Proteomics for performing the Luminex cytokine assays.

## Author Contributions

Conceptualization: L.M.G., S.P.; Methodology: L.M.G, S.O.E., A.K., S.S., V.K., S.P.; Investigation: L.M.G, J.H., H.J., T.J., A.R., A.K.; Formal analysis: L.M.G., T.J., A.K.; Visualization: L.M.G., T.J., A.R.; Funding acquisition: S.P.; Project administration: L.M.G.; Supervision: S.P. Writing – original draft: L.M.G., Writing – review & editing: L.M.G., J.H., H.J., T.J., A.K., S.O.E., S.S., V.K.,S.P.

