## Supplementary Figures for "SARS-CoV-2 S1 spike protein induces a temporal systemic immune response and promotes long-term anxiety-like behaviors"

### Supplementary Figures and Information

#### Suppl. Figure 1

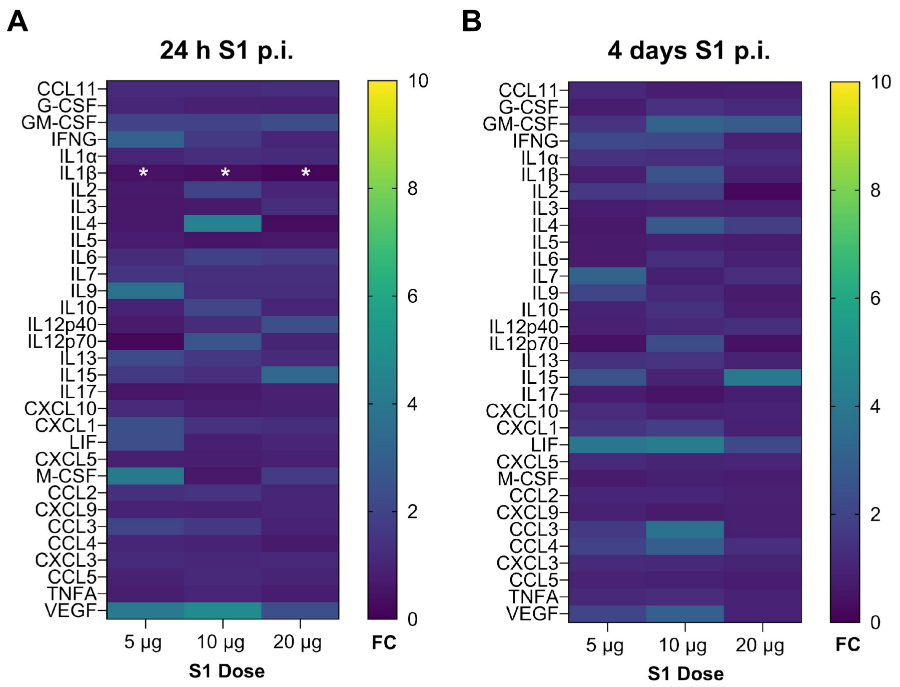


**Figure S1. SARS-CoV-2 spike S1 protein does not elicit a significant cytokine or chemokine response in wild-type mice.**

**(A, B)** Cytokine analyses of serum in control and S1-injected mice at different doses (5 µg, 10 µg and 20 µg) 24 h (**A**) and 4 days (**B**) post-inoculation (p.i.). Data shown as fold change (FC) of the concentration corresponding to the median fluorescence intensity compared with control group. We only observed a significant decrease in IL1β in S1-injected mice at 24 h p.i. Repeated measure two-way ANOVA test followed by Dunnet's *post-hoc* test: $^*P < 0.05$. n = 4 mice per group.

#### Suppl. Figure 2

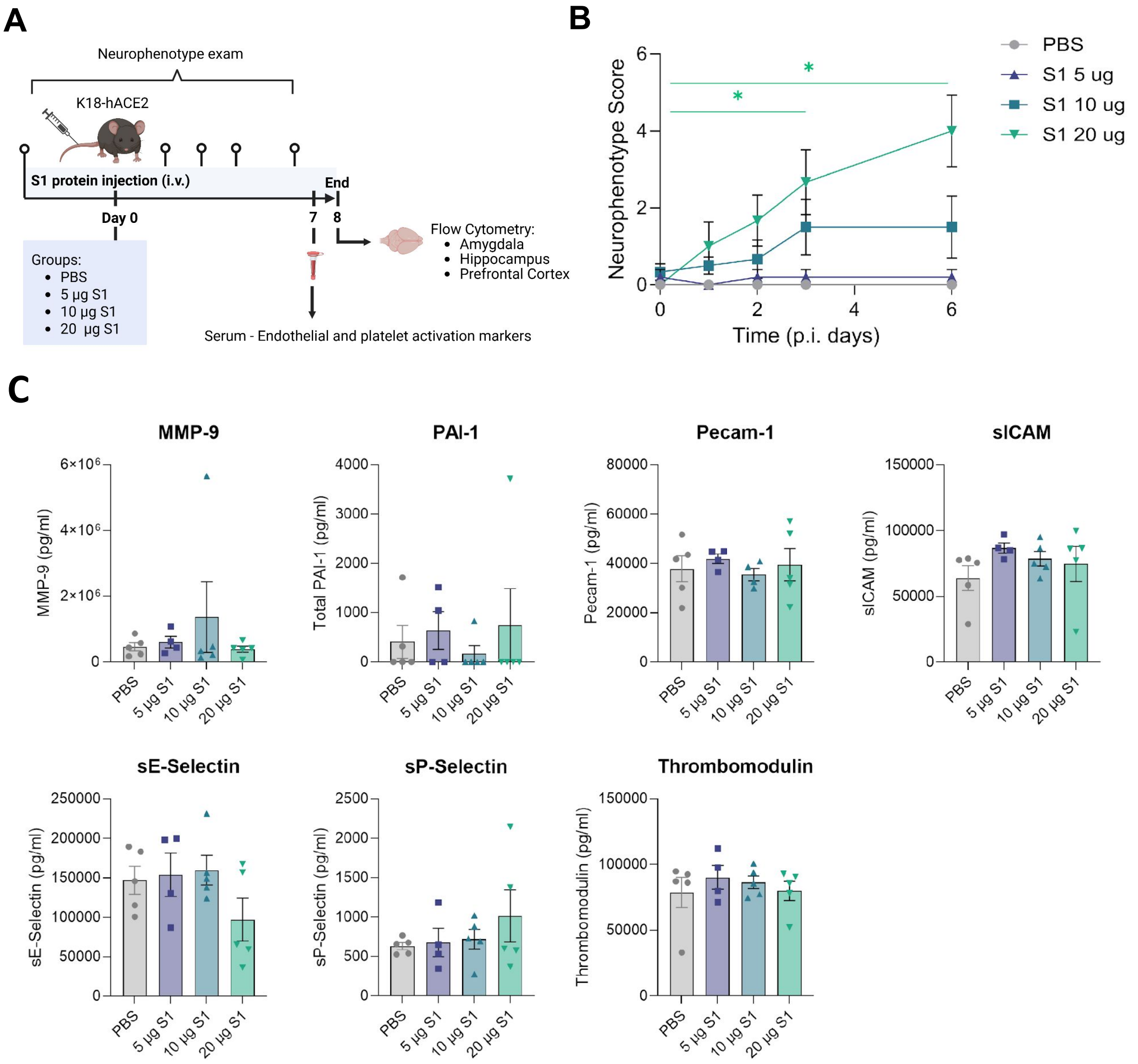


**Figure S2. SARS-CoV-2 spike S1 protein does not elicit vascular end endothelial markers elevations in K18-hACE2 mice.**

(**A**) Schematic of experimental paradigm for SARS-CoV-2 S1 protein injection (intravenous, i.v.) in K18-hACE2 mice and experimental workflow.

(**B**) Neurophenotype scores over time in control and S1-injected mice at different doses (5 µg, 10 µg and 20 µg). Repeated measure Two-way ANOVA followed by Dunnet's *post-hoc* test: *$P < 0.05$. n = 5–6 mice per group.

(**C**) Vascular and endothelial activation markers analysis of serum in control and S1-injected mice at different doses (5 µg, 10 µg and 20 µg) 7 days post-inoculation (p.i.). Data shown as the concentration (pg/ml) corresponding to the median fluorescence intensity. One-way ANOVA: no significant differences observed. n =

4–5 mice per group. Data are presented as mean ± SEM. Each dot represents an individual mouse.

### Suppl. Figure 3

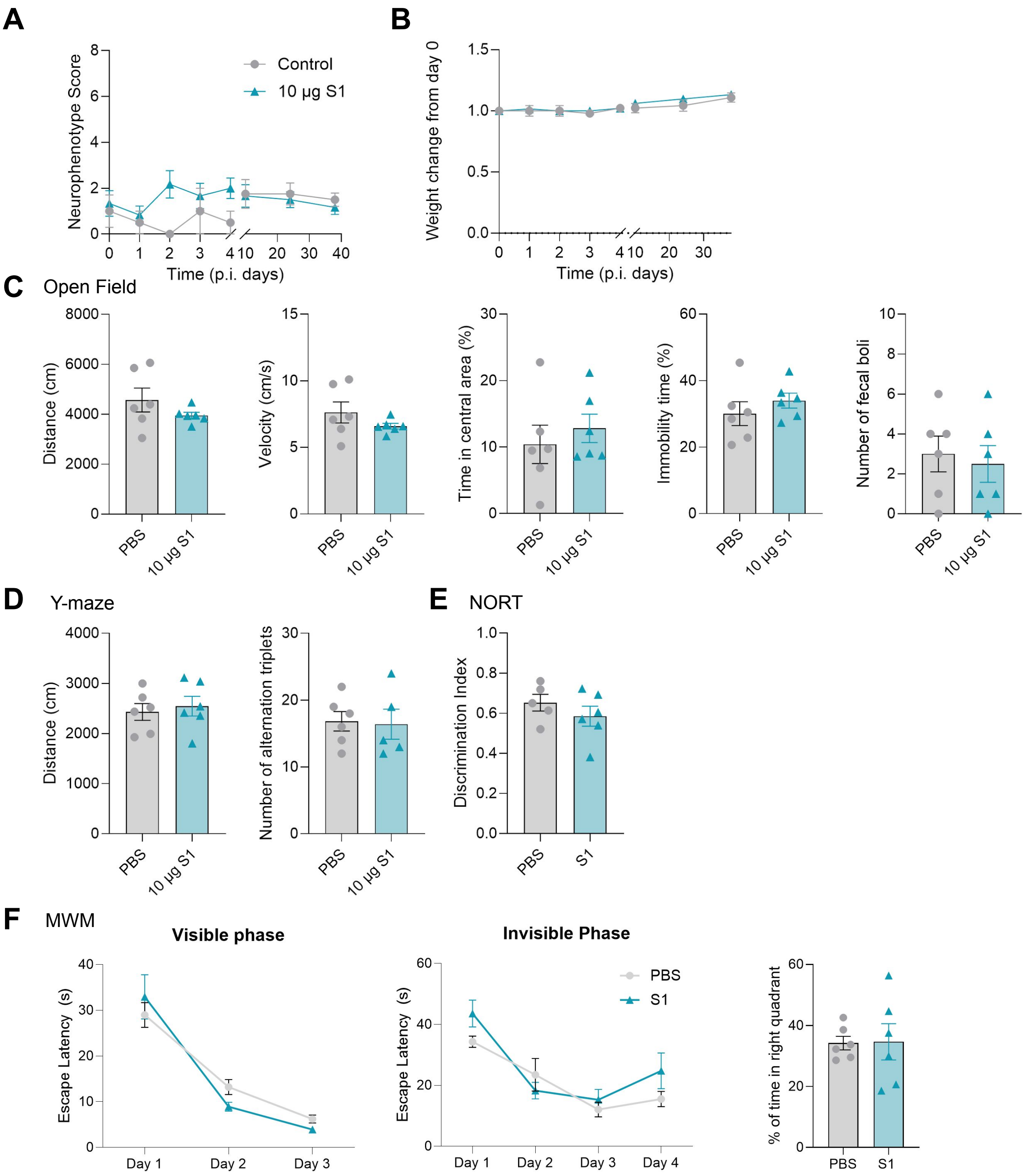

**Figure S3. SARS-CoV-2 spike S1 protein does not induce behavioral alterations in wild-type mice.**

**(A, B)** Neurophenotype scores (**A**) and body weight (change from day 0) (**B**) over time in control and S1-injected wild-type mice (10 µg dose). Repeated measures two-way ANOVA: no significant differences observed. n = 6 mice per group.

**(C)** Total distance travelled (cm), velocity (cm/s), time in central area (%), immobility time (%) and number of fecal boli in control and S1-injected wild-type mice (10 µg dose) at 2 months p.i. Unpaired t-test: no significant differences observed.

**(D)** Total distance (cm) and number of alternation triplets in control and S1-injected wild-type mice (10 µg dose) at 2 months p.i. during the Y-maze test. Unpaired t-test: no significant differences observed.

**(E)** Discrimination index—calculated as the difference in time spent exploring the familiar versus the novel object, divided by the total exploration time—in control and S1-injected wild-type mice (10 µg dose) at 2 months p.i. during the Novel Object Recognition Test (NORT). Unpaired t-test: no significant differences observed.

**(F)** Escape latency (s) during the visible and invisible phases and % of time spent in the right quadrant during the probe test of the Morris water maze (MWM), in control and S1-injected wild-type mice (10 µg dose) at 2 months p.i. Repeated measure two-way ANOVA and unpaired t-test, respectively: no significant differences observed.

Data are presented as mean ± SEM. Each dot represents an individual mouse.

#### Suppl. Figure 4

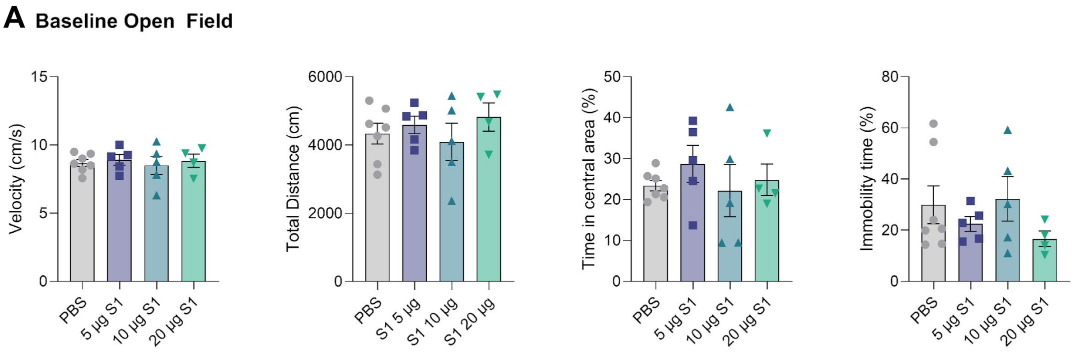


**Figure S4. No baseline open-field alterations in hACE2 mice.**

**(A)** Velocity (cm/s), total distance travelled (cm), time in central area (%), immobility time (%) and number of fecal boli in control and S1-injected hACE2 mice at different doses (5 µg, 10 µg and 20 µg) at baseline before the S1 injection (E). One-way ANOVA followed by Dunnet's *post-hoc* test: no significant differences observed.

Data are presented as mean ± SEM. Each dot represents an individual mouse. n = 4-7 mice.

#### Suppl. Figure 5

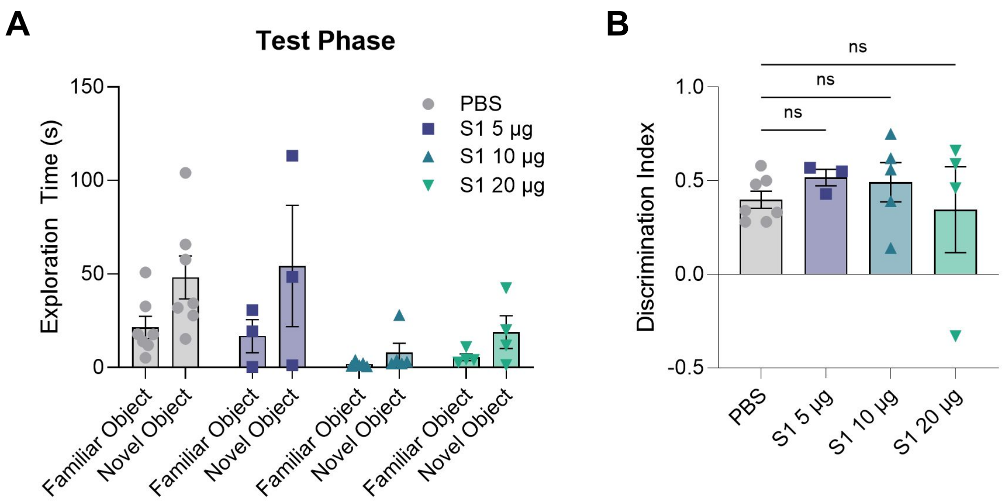


**Figure S5: Novel object recognition test results in hACE2 mice.**

**(A)** Exploration time (s) of familiar and novel objects in control and S1-injected injected hACE2 mice at different doses (5 µg, 10 µg and 20 µg) 2 months p.i. during the novel object recognition test's (NORT) test phase.

**(B)** Discrimination index—calculated as the difference in time spent exploring the familiar versus the novel object, divided by the total exploration time—in control and S1-injected injected hACE2 mice at different doses (5 µg, 10 µg and 20 µg) 2 months p.i. during the NORT. Ordinary one-way ANOVA: no significant differences observed. Note that the decrease in exploration during the familiarization phase (Figure 3) made it difficult to obtain reliable results in the subsequent test phase of the NORT.

Data are presented as mean ± SEM. Each dot represents an individual mouse. n = 4-7 mice.

### Suppl. Figure 6

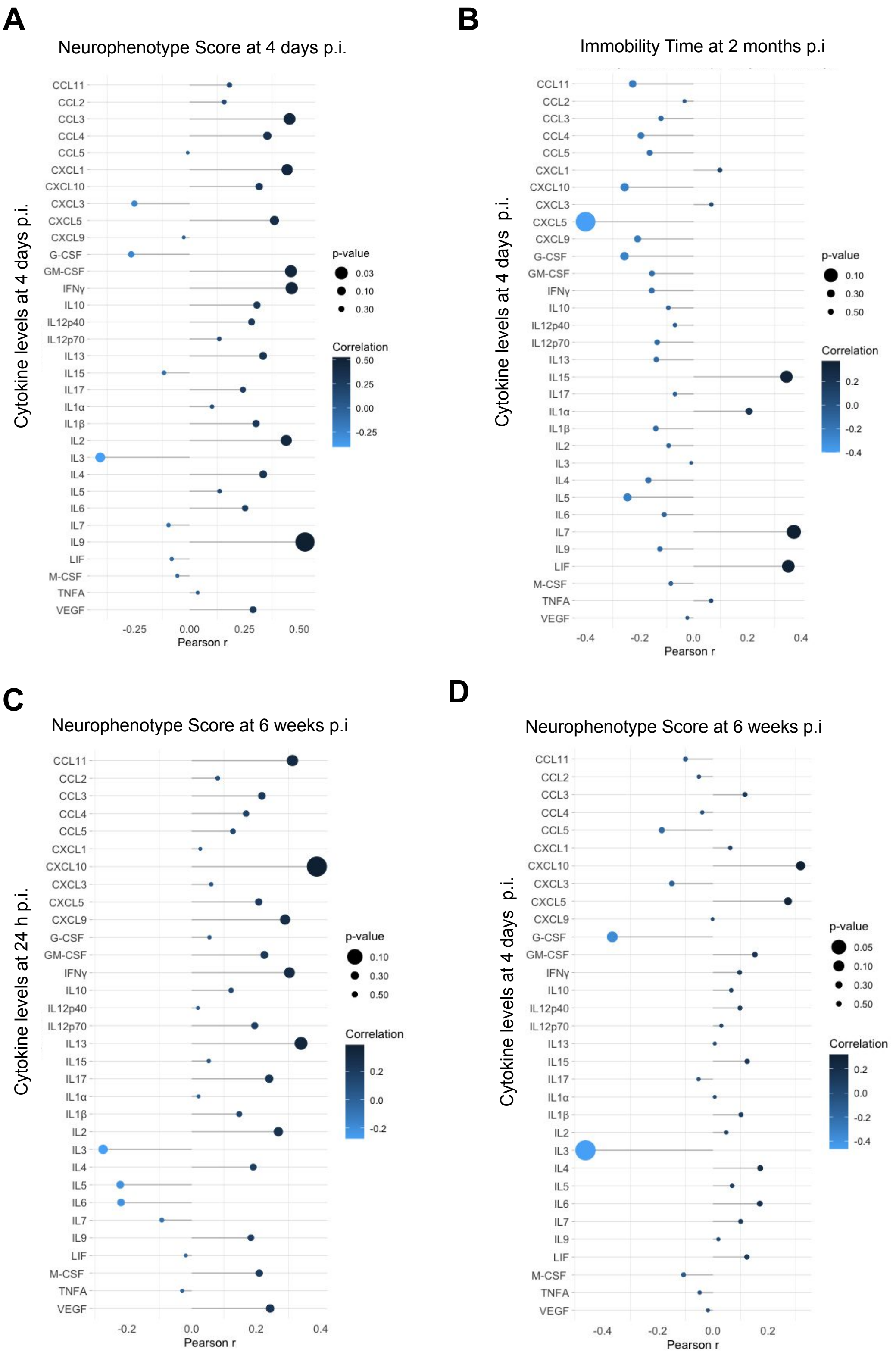


76

**Figure S6: Correlations of cytokine levels with behavioral outcomes during the acute and long-term phases.**

(**A** and **B**) Correlation of the cytokine levels at 4 days p.i. with the neurophenotype exam at 4 days p.i. (**A**) and immobility time at 2 months p.i. (**B**).

(**C, D**) Correlation of the cytokine levels at 24 h p.i. (**C**) and 4 days p.i. (**D**) and neurophenotype score at 6 weeks p.i.

Pearson's correlations. Total n = 21 mice.

### Suppl. Figure 7

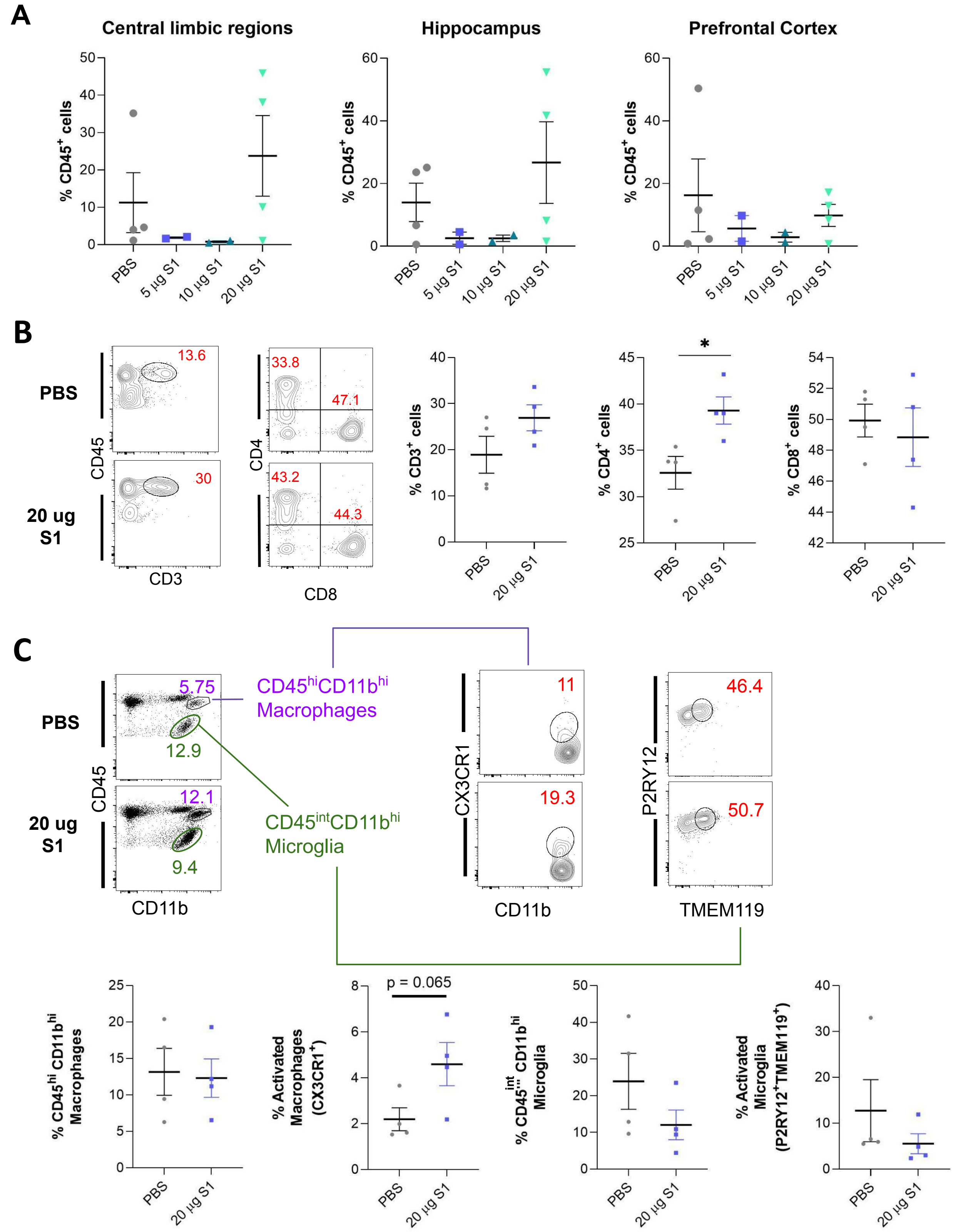

**Figure S7: SARS-CoV-2 spike S1 protein induces $CD4^+$ T cell accumulation in limbic brain regions.**

**(A)** Percentage of $CD45^+$ immune cells from live cells in central limbic regions (amygdala and hypothalamus), hippocampus and prefrontal cortex from control and S1-injected injected hACE2 mice at different doses (5 µg, 10 µg and 20 µg) 8 days p.i. Ordinary one-way ANOVA: no significant differences.

**(B – C)** Immune cell composition within $CD45^+$ cells in limbic regions of control and S1-injected hACE2 mice at different doses (5 µg, 10 µg and 20 µg) 8 days p.i. (**B**) Frequency of $CD4^+$ and $CD8^+$ T cells expressed as a percentage of total T cells. **(C)** Frequency of macrophages ($CD45^{hi}CD11b^{hi}$) and microglia ($CD45^{int}CD11b^{hi}$) within $CD45^+$ cells. Unpaired t-test: $^*P < 0.05$. n = 4 pooled samples per group.

#### Supplementary Methods

##### Flow Cytometry

After brain collection, the prefrontal cortex, hippocampus, and central limbic structures—including the amygdala and hypothalamus—were rapidly dissected on ice, placed in PBS and processed for flow cytometry (1). Briefly, excised regions of the brain were enzymatically digested with collagenase A, for 1 hour at 37 °C. Following digestion, samples were further dissociated manually and passed through a 100 µm cell strainer to obtain single-cell suspensions. Mononuclear cells were isolated using a Percoll density gradient and analyzed by flow cytometry. Immune cell types were identified using the following antibodies: CD45, CD11b, CX3CR1, P2RY12, TMEM119, CD3, CD4, and CD8. Among live singlets myeloid cells, microglia and macrophages were identified based on differential CD45 and CD11b expression. Microglia were characterized by CD45 hi CD11b lo and macrophages were designated as CD45 hi CD11b hi. The subset of cells expressing $P2RY12^+TMEM119^+$ cells within the resident microglial population were annotated as activated, whereas activated macrophages were assessed by increased expression of CX3CR1. CD169 expression status determined the inflammatory status of both microglia and macrophages. $CD45^+CD3^+$ cells were identified as total T cells. CD4 and CD8 T cell subsets were further defined as $CD3^+CD4^+$ and $CD3^+CD8^+$ populations, respectively. Following staining, cells were washed and fixed using 2% paraformaldehyde prior to acquisition on flow cytometer (BD FACSymphony A5) Data was analyzed using FlowJo software (Treestar).
